# The endothelial-specific *LINC00607* mediates endothelial angiogenic function

**DOI:** 10.1101/2022.05.09.491127

**Authors:** Frederike Boos, James A. Oo, Timothy Warwick, Stefan Günther, Judit Izquierdo Ponce, Giulia Buchmann, Tianfu Li, Sandra Seredinski, Shaza Haydar, Sepide Kashefiolasl, Andrew H. Baker, Reinier A. Boon, Marcel H. Schulz, Francis J. Miller, Ralf P. Brandes, Matthias S. Leisegang

## Abstract

Long non-coding RNAs (lncRNAs) can act as regulatory RNAs which, by altering the expression of target genes, impact on the cellular phenotype and cardiovascular disease development. Endothelial lncRNAs and their vascular functions are largely undefined. Deep RNA-Seq and FANTOM5 CAGE analysis revealed the lncRNA *LINC00607* to be highly enriched in human endothelial cells. *LINC00607* was induced in response to hypoxia, arteriosclerosis regression in non-human primates and also in response to propranolol used to induce regression of human arteriovenous malformations. siRNA knockdown or CRISPR/Cas9 knockout of *LINC00607* attenuated VEGF-A-induced angiogenic sprouting. *LINC00607* knockout in endothelial cells also integrated less into newly formed vascular networks in an *in vivo* assay in SCID mice. Overexpression of *LINC00607* in CRISPR knockout cells restored normal endothelial function. RNA- and ATAC-Seq after *LINC00607* knockout revealed changes in the transcription of endothelial gene sets linked to the endothelial phenotype and in chromatin accessibility around ERG-binding sites. Mechanistically, *LINC00607* interacted with the SWI/SNF chromatin remodeling protein BRG1. CRISPR/Cas9-mediated knockout of *BRG1* in HUVEC followed by CUT&RUN revealed that BRG1 is required to secure a stable chromatin state, mainly on ERG-binding sites. In conclusion, *LINC00607* is an endothelial-enriched lncRNA that maintains ERG target gene transcription by interacting with the chromatin remodeler BRG1.

## Introduction

Endothelial cells form the selectively permeable monolayer between the vessel and the blood. Resting endothelium provides an anti-coagulant and anti-inflammatory surface and contributes to the control of local vascular tone. It also facilitates the vascular response to inflammation, shear stress and hypoxia [12]. In response to growth factors and hypoxia, endothelial cells sprout from pre-existing vessels in the process of angiogenesis [52]. This process is physiologically important and required for wound healing [27]. However, uncontrolled angiogenesis also contributes to pathological conditions like macular degeneration and cancer [17].

Recent studies suggest that long non-coding RNAs (lncRNAs) are essential in the regulation of cardiovascular gene programs [51, 59]. LncRNAs are RNA molecules longer than 200 nucleotides in length, which may lack apparent protein-coding potential. They have independent functions as RNAs, separate from potential peptide coding abilities [51, 59]. Through different mechanisms lncRNAs impact on gene expression and therefore the cellular phenotype [59]. LncRNAs influence many aspects of cellular function among them nuclear architecture, transcription, translation and mRNA stability [64].

Transcriptional control can be exerted through interaction with or recruitment of chromatin remodeling complexes, which subsequently alter the epigenetic landscape [59]. Chromatin remodeling proteins regulate DNA accessibility by restructuring, mobilizing, and ejecting nucleosomes [9] and thereby altering the binding of transcription factors to their DNA targets [39]. One well-known multi- protein chromatin remodeling complex, the Switch/Sucrose Non-Fermentable (SWI/SNF) complex, has Brahma related gene-1 (BRG1) as one of its core catalytic subunits, whose knockout is embryonic lethal in mice [8, 28]. Several lncRNAs are known to contribute to the function of BRG1, e.g. *EVF2* directly inhibits the ATPase and chromatin remodeling activity [10], *MANTIS* stabilizes the interaction between BRG1 and BAF155 and recruits BRG1 to angiogenesis related genes [38] and *Mhrt* interacts with the helicase domain of BRG1 leading to the inhibition of chromatin target recognition by BRG1 [21]. *Xist* binding inhibits BRG1 activity and functionally antagonizes the recruitment of associated SWI/SNF complexes to the inactivated X chromosome [26]. *MALAT1* forms a complex with BRG1 and HDAC9, which inhibits the expression of contractile proteins in aortic aneurysm [43]. These examples highlight the fundamental importance of lncRNA-BRG1 interactions.

In this study, we set out to identify endothelial-enriched lncRNAs that impact on angiogenic function and may therefore have disease or therapeutic relevance. This led to the identification of the lncRNA *LINC00607*, which is highly enriched in the endothelium. *LINC00607* has been previously described as a super enhancer-derived lncRNA induced by high glucose and TNFα levels [11]. Our study revealed that *LINC00607* is induced by hypoxia and sustains endothelial gene transcription through interaction with the chromatin remodeling protein BRG1. Ultimately, *LINC00607* facilitates proper endothelial ERG-responsive gene transcription and the maintenance of the angiogenic response.

## Material and Methods

### Materials

The following chemicals and concentrations were used for stimulation: Human recombinant VEGF-A 165 (R&D, 293-VE, 30 ng/mL), DMOG (400091, Merck, 1 mM), acriflavine (A8126, Sigma-Aldrich, 10 µM), Low Density Lipoprotein from Human Plasma, oxidized (oxLDL, L34357, Thermo Fisher, 10 µg/mL), DMSO (D2650, Sigma-Aldrich), Propranolol hydrochloride (P0884, Sigma-Aldrich, 100 µM), TGF-β2 (100-35B, Peprotech, 10 ng/mL), Interleukin 1β (IL-1β, 200-01B, Peprotech, 1 ng/mL) and RNase A (EN0531, Thermo Fisher).

The following antibodies were used: β-Actin (A1978, Sigma-Aldrich) and BRG1 (ab110641, Abcam).

### Cell culture and stimulation experiments

Pooled human umbilical vein endothelial cells (HUVEC) were purchased from PromoCell (C-12203, Lot No. 405Z013, 408Z014, 416Z042, Heidelberg, Germany) and cultured at 37 °C with 5 % CO2 in a humidified incubator. Gelatin-coated dishes (356009, Corning Incorporated, USA) were used to culture the cells. Endothelial growth medium (EGM), consisting of endothelial basal medium (EBM) supplemented with human recombinant epidermal growth factor (EGF), EndoCGS-Heparin (PeloBiotech, Germany), 8 % fetal calf serum (FCS) (S0113, Biochrom, Germany), penicillin (50 U/mL) and streptomycin (50 µg/mL) (15140-122, Gibco/Lifetechnologies, USA) was used. For each experiment, at least three different batches of HUVEC from passage 3 were used.

In hypoxia experiments, cells were incubated for 24 h in a SciTive Workstation (Baker Ruskinn) at 1% O2 and 5 % CO2.

For EndMT, HUVEC were stimulated for 5 d in differentiation medium (DM) consisting of endothelial basal medium (EBM) supplemented with 8 % FCS, penicillin (50 U/mL), streptomycin (50 µg/mL), L-glutamine, TGF-β2 (10 ng/mL) and IL-1β (1 ng/mL).

### Experiments with Macaca fascicularis

Experiments on adult male Cynomolgus monkeys (*Macaca fascicularis*) were approved by the Institutional Care and Use Committee of the University of Iowa [22] and vessels were kindly provided by one of the co-authors (FJM). The vessels originated from a previous study [22], in which *Macacae fasciculari* were fed with three different diets, a normal diet, an atherosclerotic diet for 47±10 (mean ± SE) months, or an atherosclerotic diet with an additional recovery phase for 8 months. After isolation of RNA, RT-qPCR was perfomed for the orthologues of human GAPDH and *LINC00607* with *Macacae fascicularis* (*Mf*) specific primers. The following oligonucleotide sequences were used: *Mf_LINC00607*, forward 5’-CTG CAT GTC ACC GCA TAC CC-3’ and reverse 5’-TGG CTC TGC TGC TGG AGT AG-3’; *Mf_GAPDH*, forward 5’-TGC ACC ACC AAC TGC TTA GC-3’ and reverse 5’-GGC GTG GAC TGT GGT CAT GAG-3’.

### Human brain arteriovenous malformation under propranolol treatment

Patients with arteriovenous malformation (AVM) evaluated at University Hospital Frankfurt were entered into an ongoing prospective registry. The study protocol was approved by the ethical committee of the Goethe University (approval number UCT-63-2020, Frankfurt am Main, Germany). All patients with proved unruptured AVMs were included after written informed consent. Patients with arteriovenous malformation (AVM) who underwent microsurgery and had tissue available were further analyzed. We selected from our tissue bank cases of unruptured brain AVMs in patients who did not undergo pre-surgical embolization. The patients did not undergo endovascular embolization before surgical resection, and medical records did not show previous history of rupture. AVM tissue (pieces with a diameter of 0.5 cm) was cultured immediately after surgical resection in the presence of 100 μM propranolol or solvent DMSO for 72 h. Afterwards, RNA was isolated and RT-qPCR was performed.

### RNA isolation, Reverse transcription and RT-qPCR

Total RNA isolation was performed with the RNA Mini Kit (Bio&Sell) according to the manufacturers protocol and reverse transcription was performed with the SuperScript III Reverse Transcriptase (Thermo Fisher) using a combination of oligo(dT)23 and random hexamer primers (Sigma). The resulting cDNA was amplified in an AriaMX cycler (Agilent) with the ITaq Universal SYBR Green Supermix and ROX as reference dye (Bio-Rad, 1725125). Relative expression of human target genes was normalized to β-Actin, whereas for *Macaca fasciluraris* genes GAPDH was used. Expression levels were analyzed by the delta-delta Ct method with the AriaMX qPCR software (Agilent). The following oligonucleotide sequences were used: human *LINC00607*, forward 5’-CCA CCA CCA CCA TTA CTT TC-3’ and reverse 5’-AGG CTC TGT ATT CCC AAC TG-3’; human β*-Actin*, forward 5’-AAA GAC CTG TAC GCC AAC AC-3’ and reverse 5’-GTC ATA CTC CTG CTT GCT GAT-3’.

### Knockdown with siRNAs

For small interfering RNA (siRNA) treatments, HUVEC (80–90 % confluent) were transfected with GeneTrans II according to the instructions provided by MoBiTec (Göttingen, Germany). A Silencer® Select siRNA was used for siRNA-mediated knockdown of *LINC00607* (Thermo Fisher Scientific, s56342). As negative control, scrambled Stealth RNAi™ Med GC (Life technologies) was used. All siRNA experiments were performed for 48 h.

### Protein Isolation xand Western Analyses

For whole cell lysis, HUVEC were washed in Hanks solution (Applichem) and lysed with RIPA buffer (1x TBS, 1 % Desoxycholat, 1 % Triton, 0.1 % SDS, 2 mM Orthovanadat (OV), 10 nM Okadaic Acid (OA), protein-inhibitor mix (PIM), 40 µg/mL Phenylmethylsulfonylfluorid (PMSF)). After centrifugation (10 min, 16,000 xg), protein concentrations of the supernatant were determined with the Bradford assay and the extract boiled in Laemmli buffer. Equal amounts of protein were separated with SDS-PAGE. Gels were blotted onto a nitrocellulose membrane, which was blocked afterwards in Rotiblock (Carl Roth). After application of the first antibody, an infrared-fluorescent-dye-conjugated secondary antibody (Licor) was used. Signals were detected with an infrared-based laser scanning detection system (Odyssey Classic, Licor).

### LentiCRISPRv2

Guide RNAs (gRNA) targeting *LINC00607* were selected using the publicly available CRISPOR algorithm (http://crispor.tefor.net/) [20]. A dual gRNA approach consisting of gRNA-A and gRNA-B was used to facilitate the knockout of *LINC00607*. gRNA-A targeted a region downstream of the TSS and gRNA-B targeted a region upstream of the TSS. *BRG1* knockout was performed using a single gRNA approach. The gRNAs were cloned into lentiCRISPRv2 vector backbone with Esp3I (Thermo Fisher, FD0454) according to the standard protocol [55]. lentiCRISPRv2 was a gift from Feng Zhang (Addgene plasmid #52961; http://n2t.net/addgene:52961; RRID:Addgene_52961) [55]). The modification of the lentiviral CRISPR/Cas9v2 plasmid with hygromycin resistance was provided by Frank Schnütgen (Dept. of Medicine, Hematology/Oncology, University Hospital Frankfurt, Goethe University, Frankfurt, Germany).

For annealing, the following oligonucleotides were used: *LINC00607*: gRNA-A, 5’- CAC CGC ATG TGC CCC CTT TGT TGA A-3’ and 5’- AAA CTT CAA CAA AGG GGG CAC ATG C-3’, gRNA-B, 5’- CAC CGC AGT GTG TCA TGT TAT CTT G-3’ and 5’- AAA CCA AGA TAA CAT GAC ACA CTG C-3’; *BRG1*: gRNA, 5’-CAC CGC ATG CTC AGA CCA CCC AG-3’ and 5’-AAA CCT GGG TGG CTC TGA GCA TGC-3’. For *LINC00607*, gRNA-A was cloned into lentiCRISPRv2 with hygromycin resistance, gRNA-B was cloned into lentiCRISPRv2 with puromycin resistance. For *BRG1*, lentiCRISPRv2 with puromycin resistance was used. After cloning, the gRNA-containing LentiCRISPRv2 vectors were sequenced and purified. Lentivirus was produced in Lenti-X 293T cells (Takara, 632180) using Polyethylenamine (Sigma-Aldrich, 408727), psPAX2 and pVSVG (pMD2.G). pMD2.G was a gift from Didier Trono (Addgene plasmid #12259; http://n2t.net/addgene:12259; RRID:Addgene_12259). psPAX2 was a gift from Didier Trono (Addgene plasmid #12260; http://n2t.net/addgene:12260; RRID:Addgene_12260). LentiCRISPRv2-produced virus was transduced in HUVEC (p1) with polybrene transfection reagent (MerckMillipore, TR-1003-G) and for *LINC00607* knockout selection was performed with puromycin (1 μg/mL) and hygromycin (100 µg/mL) for 6 d and for *BRG1* only with puromycin (1 μg/mL) for 6 d.

Validation of the CRISPR/Cas9 knockout of *LINC00607* was performed from genomic DNA. Genomic DNA was isolated after selection. Cells were washed, collected and incubated with 500 µL lysis buffer (30 min, 56 °C, 800 rpm, 0.1 M Tris/HCl pH 8.5, 0.5 M NaCl, 0.2 % SDS, 0.05 M EDTA, 22.2 mg/mL Proteinase K). After removing cell fragments (1 min, 13.000 rpm, 4 °C), DNA was precipitated by adding the equal volume 100 % isopropanol followed by centrifugation (10 min, 13.000 rpm, 4 °C). The DNA was washed with 70 % EtOH (10 min, 13.000 rpm, 4 °C), air-dried and dissolved in TE-Buffer (10 mM Tris/HCl pH 8.0, 1 mM EDTA pH 8.0). CRISPR/Cas9 target sites were amplified by PCR with PCR Mastermix (ThermoFisher, K0171), containing forward and reverse primers (10 µM) and 100-500 ng DNA followed by agarose gel electrophoresis and ethidiumbromide staining. The following primers were used: *LINC00607* CRISPR target site, 5’-CTT CAG CCC ACT GAG TCT TG-3’ and 5’-GAG GAA CCA GCC AGA ATA GC-3’; *GAPDH*, 5’-TGG TGT CAG GTT ATG CTG GGC CAG-3’ and 5’- GTG GGA TGG GAG GGT GCT GAA CAC-3’.

### Scratch-wound migration assay

30,000 HUVEC were seeded on ImageLock 96-well plates (Essen Bioscience). Once a monolayer had formed, this was scratched the following day with a 96-pin WoundMaker tool (Essen Bioscience). EGM was then refreshed to remove dead and scraped cells. Afterwards, the cells were imaged in an Incucyte imaging system for 11 h (one image every one hour, with the “phase” image channel and 10X magnification). The Scratch Wound Cell Migration Module of the Incucyte S3 Live Cell Analysis System (Essen Bioscience) was used to monitor and analyze the cells.

### Spheroid outgrowth assay

Spheroid outgrowth assays in HUVEC were performed as described in [31]. Stimulation of spheroid outgrowth was performed with VEGF-A 165 (R&D, 293-VE, 30 ng/mL) for 16 h. Spheroids were imaged with an Axiovert135 microscope (Zeiss). The cumulative sprout length and spheroid diameter were quantified by analysis with the AxioVision software (Zeiss).

### Plasmid overexpression and Spheroid outgrowth assay

Plasmid overexpression was performed using 700,000 HUVEC and the Neon electroporation system (Invitrogen, 1,400 V, 1x 30 ms pulse) in E2 buffer for the following plasmids (7 µg per transfection): pcDNA3.1+*LINC00607* and pcDNA3.1+. Overexpression was performed for 24 h.

### RNA immunoprecipitation

To identify RNAs bound to a protein of interest, specific antibodies and protein G-coated beads were used to immunoprecipitate RNAs bound to the target protein. Cells were grown to 80 % confluence on a 10 cm plate (roughly 3 million cells) and washed once with Hanks buffer. 6 mL Hanks buffer was added to the cells on ice and irradiated with 0.150 J/cm^2^ 254 nm UV light. Cells were scraped twice in 500 µL Hanks buffer and centrifuged at 1,000 xg at 4 °C for 4 min. For nuclear protein isolation, cells were resuspended in hypotonic buffer (10 mM HEPES pH 7.6, 10 mM KCl, 0.1 mM EDTA pH 8.0, 0.1 mM EGTA pH 8.0, 1 mM DTT, 40 µg/mL PMSF) and incubated on ice for 15 min. Nonidet was added to a final concentration of 0.75 % and cells were centrifuged (1 min, 4 °C, 16000 xg). The nuclear-pellet was washed twice in hypotonic buffer, lysed in high salt buffer (20 mM HEPES pH 7.6, 400 mM NaCl, 1 mM EDTA pH 8.0, 1 mM EGTA pH 8.0, 1 mM DTT, 40 µg/mL PMSF) and centrifuged (5 min, 4 °C, 16000 xg). 10 % of the nuclear lysate was taken as input. 4 µg of antibody was pre-coupled to 30 µL protein G magnetic beads in bead wash buffer (20 mM HEPES pH 7.6, 200 mM NaCl, 1 mM EDTA pH 8.0, 1 mM EGTA pH 8.0, 1 mM DTT, 40 µg/mL PMSF) for 1 h at RT, then washed once with high salt buffer and twice with bead wash buffer. The antibody-coupled beads were added to the nuclear lysate and rotated for 1 h at 4 °C. Samples were placed on a magnetic bar and the lysate discarded. The beads were washed three times in high salt buffer (50 mM Tris-HCl, 1 M NaCl, 1 mM EDTA, 0.1 % SDS, 0.5 % Na-Deoxycholate, 1 % NP-40, 1 mM DTT, 40 µg/mL PMSF) at 4 °C for 10 min per wash. Beads were then washed twice in bead wash buffer 2 (20 mM TrisHCl, 10 mM MgCl2, 0.2 % Tween, 1 mM DTT, 40 µg/mL PMSF). For RNase A treatment, beads were placed in a buffer containing 20 mM Tris-HCl, EDTA pH 8.0 and 2 µL of RNase A (10 mg/mL) for 30 min at 37°C and then washed again in bead wash buffer. For elution of RNA, the remaining wash buffer was removed and 1 mL QIAzol (Qiagen) was added to the beads and incubated at RT for 10 min. 400 µL chloroform was added to the samples and vortexed for 10 sec followed by incubation for a further 10 min at RT. Samples were then centrifuged at 12,000 xg for 15 min at 4 °C. 500 µL of the upper aqueous phase was transferred to a new tube and 2 µL glycogen (GlycoBlue Coprecipitant, ThermoFisher, AM9515) and 500 µL isopropanol added. Samples were inverted multiple times and incubated at RT for 10 min before being centrifuged again at 12,000 xg for 10 min. The supernatant was removed and the pellet washed with 1 mL 75 % ethanol by vortexing. The pellet was centrifuged at 7,500 xg for 5 min at 4 °C, dried and resuspended in 30 µL nuclease-free water. RNA samples were reverse transcribed for qPCR as described above.

### In vivo Matrigel Plug Assay

150,000 HUVEC per plug were stained with Vybrant Dil (1:200 in 1 mL Basal Medium (EBM); Thermo Fisher, V-22885). After incubation (45 min at 37 °C, 5 min at 4 °C), cells were washed with EBM (Lonza), resuspended in EGM containing 20 % methocel (Sigma-Aldrich) and cultured in hanging drops (25 µL/drop). Harvesting of spheroids and injection of matrigel containing spheroids into SCID mice (Charles River Laboratories) was performed as described previously [34]. 21 d after injection, Isolectin GS-IB4 from *Griffonia simplicifolia*, Alexa Fluor® 647 Conjugate (I32450, Thermo Fisher) was administered intravenously and was allowed to circulate for 20 min. After transcardial perfusion of the animals, the plugs were dissected, cleaned, fixed in 4 % Paraformaldehyde (PFA) (over night) and subsequently cleared following the 3DISCO procedure [14]. Imaging was carried out with the Ultramicroscope II (UM-II, LaVision Biotec, Bielefeld) at 16x magnification (10 Zoom body + 2x Objective). Pictures were taken with a Neo 5.5 (3-tap) sCOMs Camera (Andor, Mod.No.: DC-152q-C00- FI). The ImSpectorPro Version_3.1.8 was used. Quantification of 3D Images was performed with Imaris (Bitplane Version 9.6). The surface function was used to manually delete auto fluorescence signals and artefacts. Signal background was removed using baseline subtraction. Cells were detected and counted with the Spots-Algorithm (estimated diameter = 10.0 μm; background subtraction = true; “intensity center Ch=3” above 395; Region Growing Type = Local Contrast). Lower threshold was chosen depending to the background signal. Cells were considered incorporated in the vascular network with the threshold of the “intensity Max. channel=2” above 575.

### RNA-Seq

900 ng of total RNA was used as input for SMARTer Stranded Total RNA Sample Prep Kit - HI Mammalian (Takara Bio). Sequencing was performed on the NextSeq500 instrument (Illumina) using v2 chemistry, resulting in average of 38M reads per library with 1x75bp single end setup. The resulting raw reads were assessed for quality, adapter content and duplication rates with FastQC [3]. Trimmomatic version 0.39 was employed to trim reads after a quality drop below a mean of Q20 in a window of 10 nucleotides [5]. Only reads between 30 and 150 nucleotides were cleared for further analyses. Trimmed and filtered reads were aligned versus the Ensembl human genome version hg38 (release 99) using STAR 2.7.3a with the parameter “--outFilterMismatchNoverLmax 0.1” to increase the maximum ratio of mismatches to mapped length to 10 % [13]. The number of reads aligning to genes was counted with featureCounts 1.6.5 tool from the Subread package [42]. Only reads mapping at least partially inside exons were admitted and aggregated per gene. Reads overlapping multiple genes or aligning to multiple regions were excluded. Differentially expressed genes were identified using DESeq2 version 1.26.0 [45]. Only genes with a minimum fold change of +- 1.5 (log2 +-0.59), a maximum Benjamini-Hochberg corrected p-value of 0.05, and a minimum combined mean of 5 reads were deemed to be significantly differentially expressed. The Ensemble annotation was enriched with UniProt data (release 06.06.2014) based on Ensembl gene identifiers (Activities at the Universal Protein Resource (UniProt) [1]).

### Assay for Transposase-Accessible Chromatin using sequencing (ATAC-Seq)

50,000 HUVEC were used for ATAC library preparation using lllumina Tagment DNA Enzyme and Buffer Kit (Illumina). The cell pellet was resuspended in 50 µL of the lysis/transposition reaction mix (25 µL TD-Buffer, 2.5 µL Nextera Tn5 Transposase, 0.5 µL 10 % NP-40 and 32 µL H2O) and incubated at 37 °C for 30 min followed by immediate purification of DNA fragments with the MinElute PCR Purification Kit (Qiagen). Amplification of Library and Indexing was performed as described elsewhere [7]. Libraries were mixed in equimolar ratios and sequenced on NextSeq500 platform using V2 chemistry. Trimmomatic version 0.39 was employed to trim raw reads after a quality drop below a mean of Q20 in a window of 5 nt [5]. Only reads above 15 nt were cleared for further analyses. These were mapped versus the hg38 version (emsambl release 101) of the human genome with STAR 2.7.7a [13] using only unique alignments to exclude reads with uncertain arrangement. Reads were further deduplicated using Picard 2.21.7 [6] to avoid PCR artefacts leading to multiple copies of the same original fragment. The Macs2 peak caller version 2.1.1 was employed to accommodate for the range of peak widths typically expected for ATAC-Seq [66]. Minimum qvalue was set to -4 and FDR was changed to 0.0001. Peaks overlapping ENCODE blacklisted regions (known misassemblies, satellite repeats) were excluded. In order to be able to compare peaks in different samples, the resulting lists of significant peaks were overlapped and unified to represent identical regions. The counts per unified peak per sample were computed with BigWigAverageOverBed [30]. Raw counts for unified peaks were submitted to DESeq2 (version 1.20.0) for normalization [45]. Peaks were annotated with the promoter of the nearest gene in range (TSS +- 5000 nt) based on reference data of GENCODE vM15. Peaks were deemed to have significantly different counts between conditions at an average score of 20, and a log2 transformed fold change of <-0.59 or >0.59.

### RNA Fluorescence in-situ hybridization

Cells grown on gelatin-coated 8-well µ-Slides (ibidi) were fixed in 4 % PFA (in PBS, 10 min, at RT) and washed 3 times with PBS. Cells were permeabilized in 0.5 % Triton X-100 (in PBS, 5 mM vanadyl complex (VRC, NEB)) on ice for 10 min and washed 3 times with PBS. Prior to hybridization, cells were rinsed once in 2xSSC. Hybridization was performed over night at 37 °C in hybridization buffer (10 % dextran sulfate, 50 % formamide, 2xSSC, 400 µg E.coli tRNA, 0.02 % RNase-free bovine serum albumin, 2 mmol/L VRC) and 10 nmol/L 5’TYE-665 labelled locked nucleic acid (LNA) detection probe (Qiagen). Custom LNA detection probes targeting *LINC00607* were designed with the Qiagen GeneGlobe Custom LNA design tool and had the following sequences: 5’-AGG AGC TGA GAT GCA CAT ACT-3’. The cells were washed 4 times for 15 min in buffer containing 2xSSC and 50 % formamide and were counterstained with DAPI (in PBS). Images were captured with a laser confocal microscope LSM800 (Zeiss, Germany) and analyzed with ZEN lite software (Zeiss, Germany).

### BRG1 CUT&RUN

BRG1 Cleavage Under Targets & Release Using Nuclease (CUT&RUN), a method established by Skene and Henikoff in 2017 [58], was performed similarly as described in the EpiCypher CUT&RUN Protocol v2.0, but with minor modifications for the cell type and antibody used. Briefly, 500,000 NTC or *BRG1* knockout HUVEC were washed with wash buffer (20 mM HEPES pH 7.9, 150 mM NaCl, 500 nM spermidine, 1X Roche Protein Inhibitor Cocktail) at RT. Cells were resuspended in wash buffer and 10 µL BioMag®Plus Concanavalin A (ConA) beads (Polysciences, 86057-3) were added for 10 min at RT. Beads were separated on a magnetic rack and washed once before being resuspended in 100 µL antibody buffer (wash buffer, 0.25 % Digitonin and 2 mM EDTA) and 1 µL BRG1 antibody (Abcam, ab110641). Beads were incubated with the antibody over night with gentle shaking at 4 °C. The next day, beads were washed twice with 200 µL 0.25 % Digitonin wash buffer and resuspended in Digitonin wash buffer containing 2 µL CUTANA™ pAG-MNase (15-1016, EpiCypher, 15-1016) and incubated on ice for 30 min. Samples were washed twice and then resuspended in 100 µL Digitonin wash buffer containing 2 µL CaCl2 at a final concentration of 100 mM and incubated for 2 h at 4 °C with gentle shaking. 33 µL of 2X “stop solution” (340 mM NaCl, 20 mM EDTA, 4 mM EGTA, 0.25 % Digitonin, 100 µg/mL RNase A, 50 µg/mL Glycoblue) was added to the beads and incubated at 37 °C for 10 min. Samples were placed on a magnetic rack and the supernatant removed and kept for DNA purification. Briefly, 5X volume of binding buffer (20 mM HEPES pH 7.9, 20 mM KCl, 1 mM CaCl2, 1 mM MnCl2) was added to the samples and the pH adjusted with sodium acetate before being transferred to a purification column (ActiveMotif, 58002) and centrifuged at 11,000 xg for 30 sec. The column was then washed with 750 µL wash buffer and dried by centrifugation for 2 min. DNA was eluted with 25 µL elution buffer and the DNA concentration measured with a Qubit 3.0 Fluorometer (Life Technologies).

### Library preparation and sequencing of CUT&RUN samples

DNA libraries were prepared according to the manufacturer’s protocol (NEBNext® Ultra II, NEB) with some minor adjustments for CUT&RUN samples. Briefly, samples were brought to 50 µL with 0.1X TE buffer and DNA end preparation performed as instructed but with incubation at 20 °C for 20 min and then 58 °C for 45 min. Adaptor ligation was performed with a 1:10 dilution of adaptor (NEB, E6440S). For DNA purification, 0.9X Volume AMPure XP beads (Beckman Coulter, A63881) was added to the samples and incubated for 5 min at RT. Beads were washed twice with 200 µL 80 % ethanol and DNA eluted with 17 µL 0.1X TE buffer for 2 min at RT. PCR amplification of the eluted DNA was performed as described in the manufacturer’s protocol but with the addition of 2.5 µL Evagreen (20X) for visualization of the amplification curves on an AriaMx Real-time PCR system (Agilent). The denaturation and annealing/extension steps of the PCR amplification were performed for around 12 cycles and stopped before the curves plateaued. A cleanup of the PCR reaction was performed twice with 1.1X Ampure beads and eluted each time in 33 µL 0.1X TE buffer. DNA concentrations were measured with a Qubit (Thermo Fisher) and size distributions measured on a Bioanalyzer (Agilent).

Sequencing was performed on the NextSeq1000/2000 (Illumina). The resulting raw reads were assessed for quality, adapter content and duplication rates with FastQC [3]. Trim Galore! [15] was used to trim reads before alignment to the Ensembl human genome version hg38 (ensembl release 104) using Bowtie2 [35, 36]. Duplicate reads were removed with rmdup [40] and coverage tracks generated with bamCoverage (deepTools Version 3.5.1) [53]. ComputeMatrix [53] and plotHeatmap (deepTools Version 3.5.1) were used on the coverage tracks to generate heatmaps of BRG1 binding across the genome. Peaks were called on the aligned data using MACS2 [16] and annotatePeaks (HOMER) [23] was used to identify the nearest genes to called peaks.

### Publicly available datasets

The following RNA-Seq datasets used in this study originated from NCBI GEO: HUVEC treated with normoxia or hypoxia (GSE70330)[18]; ACF treatments of HUVEC under normoxia (GSE176555) or hypoxia (GSE186297)[57]; EndMT treatments of HUVEC and PAEC (GSE118446)[48].

FANTOM5 CAGE and ENCODE expression data was obtained from the FANTOM5 website and was published elsewhere[19, 44, 50].

Publically available HUVEC ERG ChIP-sequencing data (GSE124891) was downloaded from the Gene Expression Omnibus (GEO) [29].

### ERG ChIP-Seq data analysis

FASTQ files were trimmed with Trim Galore! [15] and aligned to the Ensembl human genome version hg38 (ensembl release 104) using Bowtie2 [35, 36]. Duplicate reads were removed with rmdup [40]. Peaks were called on the aligned data using MACS2 [16] and annotatePeaks (HOMER) [23] used to identify the nearest genes to called peaks.

### Use of FANTOM5 CAGE ENCODE data for promoter and expression analysis of LINC00607

The promoter of *LINC00607* was defined as nucleotide sequence with a length of 1000 nt, starting from a prominent FANTOM5 CAGE region having multiple peaks in close vicinity (approx. 30 nt) going in upstream direction for 970 nt (hg38 chr2:215,848,858-215,849,857). Promoter analysis was performed with filters for the indicated transcription factors with the MoLoTool (https://molotool.autosome.org/), an interactive web application suitable to identify DNA sequences for transcription factor binding sites (TFBS) with position weight matrices from the HOCOMOCO database [33].

To compare the individual lncRNA expression towards all other cell types or tissues, each cell type-specific signal obtained with FANTOM5 CAGE (or ENCODE) [19, 44, 50] was divided through the mean signal observed in all cell types or tissues and plotted.

### Gene-Set Enrichment Analysis

GSEA (Gene-Set Enrichment Analysis, http://software.broadinstitute.org/gsea/index.jsp)[49, 60] was performed based on the RNA-seq data in order to identify gene sets that were significantly enriched from genes differently expressed between the NTC control and *LINC00607* knockout. 1000 permutations were performed and gene sets were considered statistically enriched with a nominal P <0.05.

### Differential ATAC-sequencing analysis and intersection with gene-linked regulatory elements

Alignment files arising from ATAC-sequencing data analysis detailed above were subjected to replicate-based differential peak calling using *THOR* (v0.13.1) [2], which employs a hidden Markov model-based approach to identify differentially accessible regions of chromatin between conditions. Differential peaks were those with reported adjusted *p*-values less than 0.01. Differential peaks were subsequently intersected with regulatory elements from *EpiRegio* (v1.0.0) [4], a collection of regulatory elements and their associated genes. Genes whose expression is dependent on differentially accessible regulatory elements were subjected to pathway enrichment analysis using the *ReactomePA* (v1.36.0) [65] package for *R*. Subsequently, differential accessibility of regulatory elements linked to genes differentially expressed in RNA-seq could be quantified for different gene sets, and displayed graphically with *ggplot2* (v3.3.5) [63]. Motif enrichment analysis of differential ATAC-sequencing peaks was performed using *HOMER* (v4.11.1) [23] by providing sequences underlying the peaks, and otherwise the default parameters.

### Data availability

The RNA-Seq and ATAC-Seq datasets have been deposited in private status at NCBI GEO with the accession number GSE199878.

BRG1 CUT&RUN datasets have been deposited in private status at NCBI GEO with the accession number GSE201824.

### Statistics

Unless otherwise indicated, data are given as means ± standard error of mean (SEM). Calculations were performed with Prism 8.0 or BiAS.10.12. The latter was also used to test for normal distribution and similarity of variance. In case of multiple testing, Bonferroni correction was applied. For multiple group comparisons ANOVA followed by post hoc testing was performed. Individual statistics of dependent samples were performed by paired t-test, of unpaired samples by unpaired t-test and if not normally distributed by Mann-Whitney test. P values of <0.05 was considered as significant. Unless otherwise indicated, n indicates the number of individual experiments.

## Results

### LINC00607 is a highly endothelial-enriched lncRNA induced by hypoxia

A screen for the top-expressed endothelial lncRNAs in the FANTOM5 CAGE (Cap Analysis of Gene Expression)-ENCODE database revealed that *LINC00607* is one of the most endothelial-enriched lncRNAs (**Fig. 1A**). Particularly high levels of *LINC00607* were observed in aortic, venous, lymphatic, thoracic and arterial ECs (**Fig. 1B**). Additionally, FANTOM5 CAGE-ENCODE cell-type expression data showed *LINC00607* to be predominantly localized in the nucleus (**Fig. 1A**), which was confirmed by RNA-fluorescence *in situ* hybridization (RNA-FISH) in HUVEC (**Fig. 1C**). RT-qPCR after reverse transcription with random or oligodT oligonucleotides revealed that *LINC00607* has a poly-A tail (**Fig. S1A**).

**Figure 1:**
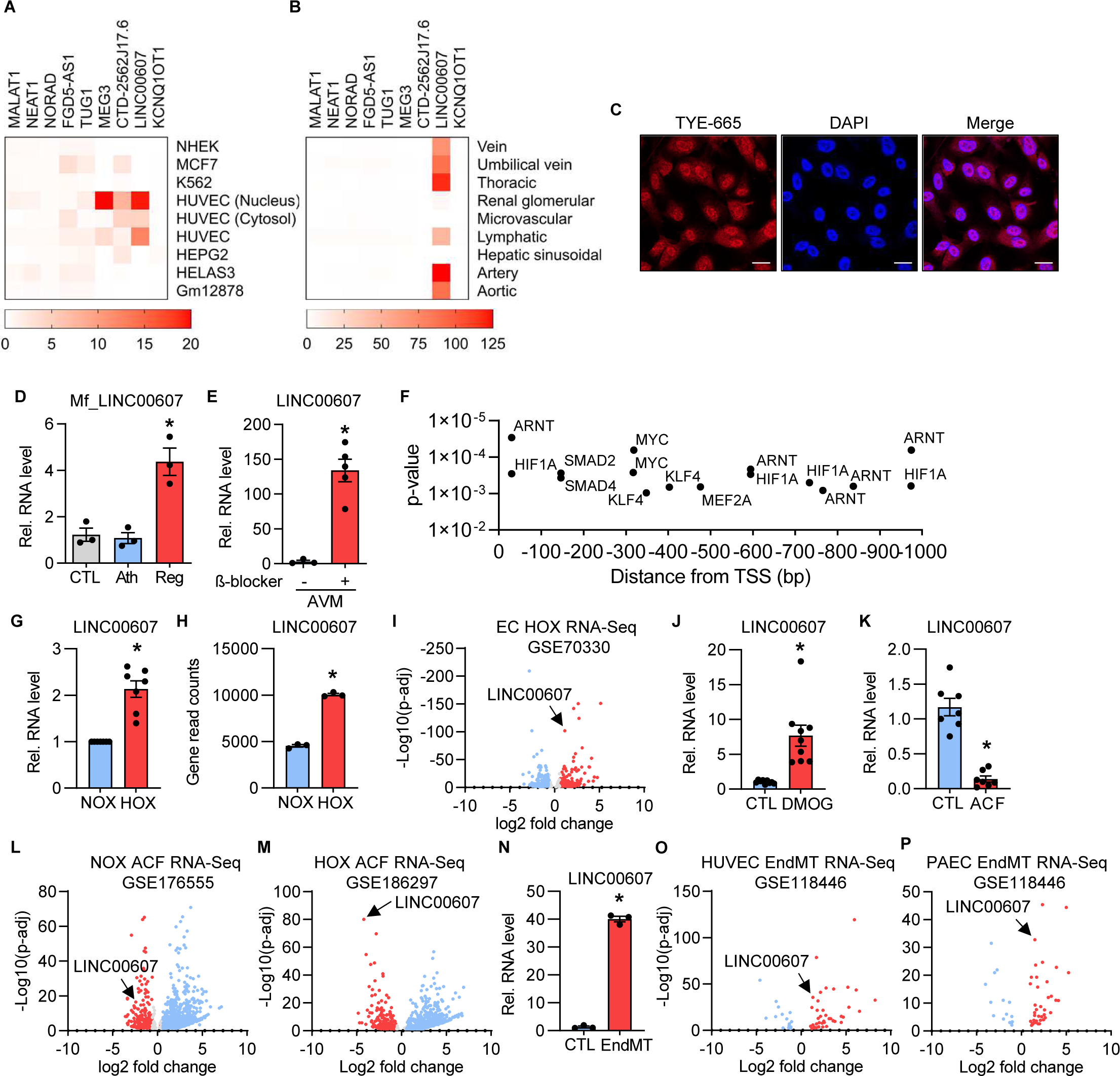
*LINC00607* is an EC-enriched lncRNA upregulated during hypoxia and EndMT. **A**, FANTOM5 CAGE-ENCODE expression of the 10 highest endothelial expressed lncRNAs across different cell lines. To compare the individual lncRNA expression towards all other cell types, each cell type-specific signal was divided through the mean signal observed in all cell types. **B**, FANTOM5 CAGE expression of the 10 highest expressed endothelial lncRNAs across different endothelial tissues. Calculation was performed as in A, but here the signals for cell tissues were used. **C**, RNA-FISH of *LINC00607* in HUVEC. *LINC00607* is labelled with a 5’TYE-665 probe, DAPI is used to stain the nuclei. Scale bar indicates 20 µm. **D**, RT-qPCR of the *LINC00607* homologue in monkey vessels originating from *Macaca fascicularis* (Mf) treated either with a normal diet (CTL), a high fat diet (Ath) or with a high-fat diet and a subsequent recovery phase (Reg). n=3. One-way ANOVA with Bonferroni post hoc test. **E**, RT-qPCR of *LINC00607* in human arteriovenous malformations (AVM) treated with and without the β- blocker propanolol for 72 h. n=5, Mann-Whitney U test. **F**, Promoter analysis of *LINC00607*. A region starting from the *LINC00607* transcriptional start site (TSS) and 1000 base pairs (bp) upstream was analyzed with MoLoTool and plotted according to p-value. **G**, Relative *LINC00607* expression in HUVEC treated with normoxia (NOX; 20% O_2_) or hypoxia (HOX; 1% O_2_), n=7. Paired t-test. **H**, *LINC00607* gene read counts in HUVEC cultured under normoxic and hypoxic conditions, n=3. Unpaired t-test. **I,** Volcano plot of log2 fold changes of lncRNAs expressed in hypoxia versus normoxia. **J**, Relative expression of *LINC00607* in HUVEC after stimulation with DMOG. DMSO served as control (CTL), n=9, Mann-Whitney U Test. **K**, Relative expression of *LINC00607* in HUVEC after stimulation with acriflavine (ACF), n=7, Mann-Whitney U Test. **L-M**, Volcano plot of log2 fold changes of lncRNAs in HUVEC treated with acriflavine (ACF) cultured under normoxia (k) or hypoxia (l). **N**, Relative expression of *LINC00607* in HUVEC under basal (CTL) or Endothelial-to-mesenchymal transition (EndMT) conditions. n=3, Unpaired t-test. **O-P**, Volcano plot of log2 fold changes of lncRNAs after EndoMT versus unstimulated control in HUVEC (O) or pulmonary arterial endothelial cells (PAEC) (P). Error bars are defined as mean +/- SEM. *p<0.05. p-adj, p-adjusted value.

Importantly, *LINC00607* expression was altered in various cardiovascular diseases. The corresponding orthologue of *LINC00607* (**Fig. S1B**) was strongly induced in *Macaca fascicularis* samples undergoing atherosclerosis regression after a high fat diet (**Fig. 1D**). Furthermore, *LINC00607* expression was increased in response to propanolol treatment of human arteriovenous malformation explants (**Fig. 1E**).

We next searched for potential gene regulatory mechanisms responsible for controlling *LINC00607* expression. An analysis of the promoter region, defined here as the FANTOM5 CAGE transcription start site signal to 1000 nucleotides (nt) upstream, revealed binding motifs for multiple transcription factors. In particular, ARNT (also known as Hypoxia Inducible Factor 1 Beta) and HIF1A (Hypoxia Inducible Factor 1 Alpha) were identified multiple times and in close proximity to the transcription start site, indicative of transcriptional regulation by hypoxia (**Fig. 1F**). Indeed, *LINC00607* expression was significantly increased when HUVEC were cultured under hypoxic conditions (1% oxygen) (**Fig. 1G**). A publicly available RNA-Seq dataset containing hypoxia-stimulated HUVEC [18] confirmed this finding (**Fig. 1H**); in fact, *LINC00607* was among the top upregulated lncRNAs in this dataset (**Fig. 1I**). Interestingly, stimulation of HUVEC with oxLDL and DMOG, the latter of which is known to stabilize HIF1α under both hypoxic and normoxic conditions [25], increased *LINC00607* expression (**Fig. 1J, S1C**). Conversely, the DNA topoisomerase and HIF-inhibitor acriflavine (ACF) [57] led to a decrease in *LINC00607* expression, which was exacerbated under hypoxia (**Fig. 1K-M**).

In addition to HIF binding sites, the promoter analysis of *LINC00607* yielded SMAD binding motifs. To test their relevance for *LINC00607* expression, HUVEC were stimulated with TGF-ß2 and IL-1ß to induce endothelial to mesenchymal transition (EndMT), a process in which SMADs play a central role [32]. Indeed, EndMT strongly increased the expression of *LINC00607* (**Fig. 1N**) and similar findings could be retrieved from publicly available RNA-Seq datasets [47] of HUVEC (**Fig. 1O**) and pulmonary arterial endothelial cells (PAEC) (**Fig. 1P**).

These data indicate that *LINC00607* is an endothelial-enriched lncRNA induced by transcription factors that are central in hypoxic and EndMT signalling.

### LINC00607 promotes sprouting, proliferation and vascularization

In order to study the functional relevance of *LINC00607* in endothelial cells, spheroid outgrowth assays were performed. In this assay, knockdown of *LINC00607* with siRNA (**Fig. 2A**) suppressed sprouting in response to VEGF-A (**Fig. 2B-D**). Next, a *LINC00607* knockout in HUVEC was achieved by CRISPR/Cas9- mediated removal of the transcriptional start site of *LINC00607* **(Fig. S1D)**. Successful knockout was confirmed on the levels of both the DNA (**Fig. 2E**) and RNA (**Fig. 2F&G**). As with siRNA-mediated knockdown, knockout of *LINC00607* inhibited VEGF-A-induced sprouting (**Fig. 2H-J**). As a second functional assay, scratch wound experiments were performed to determine migratory capacity but also proliferation. Also in this assay, the loss of *LINC00607* negatively affected endothelial function (**Fig. 2K&L**). In order to study the mechanistic function of *LINC00607*, the RNA was overexpressed in knockout cells. In the case of *cis*-action, i.e. local action of the RNA at the transcription site or a general transcriptional importance of the gene locus, such a rescue experiment should not restore function. However, transfection of *LINC00607* into *LINC00607* knockout cells restored a normal angiogenic response to VEGF-A (**Fig. 2M&N**). This suggests that the RNA itself mediates the observed functional effects by acting in *trans*.

**Figure 2:**
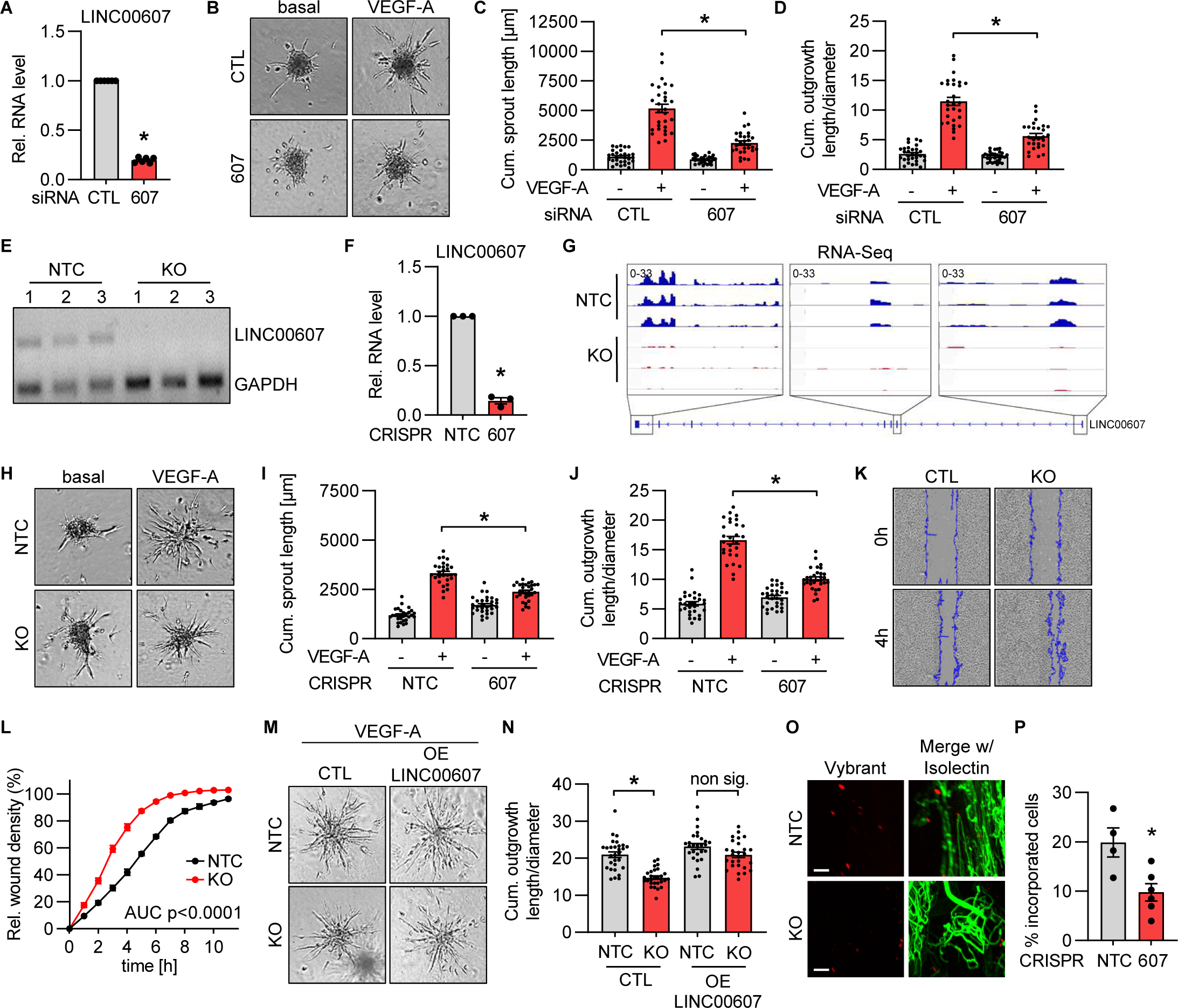
CRISPR/Cas9 KO and siRNA-knockdown reveal that *LINC00607* is important for normal EC function. **A**, RT-qPCR of *LINC00607* after siRNA-based knockdown for 48 h of *LINC00607* (607). Scrambled siRNA served as negative control (CTL). n=6. Paired t-test. **B**, Spheroid outgrowth assay after siRNA-based knockdown of *LINC00607* (607). Scrambled siRNA served as negative control (CTL). Cells treated with or without VEGF-A (16 h) are shown. **C**, Quantification of cumulative sprout length from spheroid outgrowth assay shown in Fig. 2B, n=28-30, One-way ANOVA with Bonferroni post hoc test. **D**, Quantification of the ratio of cumulative outgrowth length and respective spheroid diameter from the spheroid outgrowth assay shown in Fig. 2B; n=28-30, One-way ANOVA with Bonferroni post hoc test. **E**, PCR of Genomic DNA after lentiviral CRISPR/Cas9-mediated knockout (KO) of *LINC00607*. Three different batches (1-3) of HUVEC are shown. Non-targeting control gRNAs (NTC) served as negative control. GAPDH served as housekeeping gene. **F**, RT-qPCR of *LINC00607* after CRISPR/Cas9-mediated knockout (KO) and control (NTC), n=3. Paired t-test. **G**, IGV genome tracks of RNA-Seq of the *LINC00607* locus in HUVEC with or without CRISPR/Cas9-mediated knockout of *LINC00607* (KO) and control (NTC). **H**, Spheroid outgrowth assay with *LINC00607* knockout (KO) and control (CTL) in HUVEC. NTC served as negative control. Cells treated with or without VEGF-A are shown. **I**, Quantification of cumulative sprout length from spheroid outgrowth assay shown in Fig. 2H, n=28-30, One-way ANOVA with Bonferroni post hoc test. **J**, Quantification of the ratio of cumulative outgrowth length and respective spheroid diameter from spheroid outgrowth assay shown in Fig. 2H; n=28-30, One-way ANOVA with Bonferroni post hoc test. **K**, Scratch wound assay of *LINC00607* KO and NTC control cells (CTL). Representative images after 0 h and 4 h after scratch (blue line) are shown. **L**, Quantification of relative wound closure in *LINC00607* KO and control (NTC) in HUVEC. n=3, area under the curve (AUC) p<0.0001 (two-tailed t-test). **M**, Spheroid outgrowth assay of HUVEC after CRISPR/Cas9-mediated *LINC00607* KO or non target control (NTC). Cells treated under VEGF-A conditions for 16 h with/ without *LINC00607* overexpression (OE) are shown. Empty vector transfection served as control (CTL). **N,** Quantification of the ratio of cumulative outgrowth length and respective spheroid diameter from spheroid outgrowth assay shown in Fig. 2M. n=26-29, One-way ANOVA with Bonferroni post hoc test. **O**, *LINC00607* KO and control cells (NTC) after *in vivo* matrigel plug assay in SCID mice. HUVEC were embedded in matrigel, stained with Vybrant dil (red) and injected. Isolectin GS-b4 Alexa 647 conjugated stained vessels (green). Images were taken by light sheet microscopy 21 d after injection. Scale bar indicates 100 µm. Representative pictures are shown. **P**, Quantification of cells per plug integrated into the newly formed vascular network shown in 2M. n=5-6. Mann Whitney U test. Error bars are defined as mean +/- SEM. *p<0.05.

Collectively, these data demonstrate that loss of *LINC00607* limits endothelial angiogenic capacity. As *LINC00607* is not conserved to mice, its physiological importance was studied by assessing the capacity of HUVEC to integrate into the vascular network of matrigels when injected in SCID-mice. Importantly, in this *in vivo* assay, knockout of *LINC00607* significantly decreased the capacity of HUVEC to be integrated into the murine vascular network (**Fig. 2O&P**). These data demonstrate that *LINC00607* acts *in trans* as a pro-angiogenic lncRNA.

### LINC00607 maintains transcription of genes involved in VEGF-signalling

To identify how *LINC00607* impacts on angiogenic function, gene expression was determined by RNA- Seq with and without LentiCRISPR-mediated knockout of *LINC00607* in HUVEC. Deletion of *LINC00607* markedly impacted endothelial gene expression (**Fig. 3A-C, S2A-C, Table S1**), with a greater tendency to decrease rather than increase the expression of protein-coding and non-coding RNAs (**Fig. S2D&E)**. Due to the observed angiogenic defects, a Gene Set Enrichment Analysis (GSEA) was performed for the VEGF-signaling pathway. GSEA revealed a strong association of differentially expressed genes within the VEGF-signaling pathway after CRISPR/Cas9-mediated knockout of the lncRNA (**Fig. 3D**). This GSEA result was associated with numerous VEGF-signaling genes that were mainly downregulated upon knockout of *LINC00607* (**Fig. 3E-G**).

**Figure 3:**
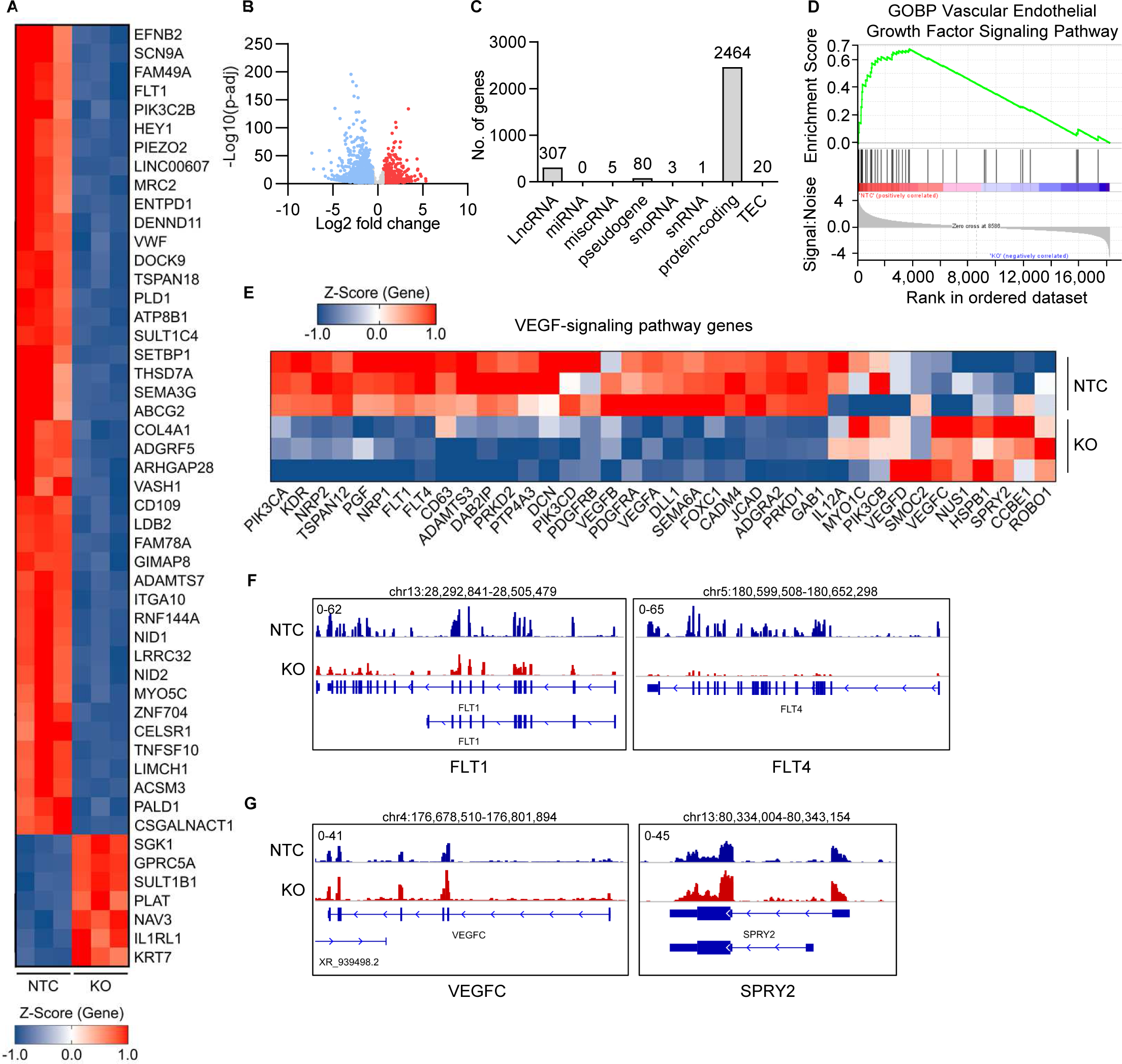
RNA- and ATAC-Seq reveal that *LINC00607* maintains endothelial gene expression. **A**, Heatmap of the top 50 differentially expressed genes as determined by RNA-Seq with (KO) or without (NTC) CRISPR/Cas9-mediated knockout of *LINC00607*. Three different batches of HUVEC are shown. Genes shown have a padj<0.05, and a log2 Fold Change greater than ±0.585. Z-score represents up- (red, positive value) or down-regulated (blue, negative values) genes. **B**, Volcano plot of RNA-Seq showing the log2 fold changes (KO vs. NTC) of all genes expressed against their p-adjusted value (p- adj). **C**, Numbers of genes from different gene classes significantly altered by *LINC00607* KO vs. NTC HUVEC determined by RNA-Seq. **D**, Gene Set Enrichment Analysis (GSEA) of significantly altered genes showing an enrichment score and signal to noise ratio for the Gene Ontology biological process (GOBP) Vascular Endothelial Growth Factor Signaling Pathway. **E**, Heat map of VEGF-signaling pathway genes and their expression differences after RNA-Seq. Z-score represents up- (red, positive value) or down- regulated (blue, negative values) genes. **F**, Examples of significantly downregulated genes after *LINC00607* knockout. IGV genome tracks of the *FLT1* and *FLT4* locus. Shown are RNA-Seq reads in *LINC00607* knockout (KO, red) and control (NTC, blue). Tracks of three replicates are overlaid. **G**, Examples of significantly upregulated genes after *LINC00607* knockout. IGV genome tracks of the *VEGFC* and *SPRY2* locus. Shown are RNA-Seq reads in *LINC00607* knockout (KO, red) and control (NTC, blue). Tracks of three replicates are overlaid. Genomic coordinates correspond to hg38.

### LINC00607 depletion reduces the accessibility of ETS transcription factor binding sites

In order to determine whether the effects of *LINC00607* loss of function and differential gene expression were a consequence of altered chromatin accessibility, an assay for transposase-accessible chromatin with sequencing (ATAC-Seq) was performed (**Fig. S2F, Table S2**). Comparison of ATAC-Seq and RNA-Seq for the multiple VEGF-signaling genes revealed a similar effect of *LINC00607* knockout on chromatin accessibility of gene-linked enhancers (as annotated by EpiRegio [4]) and gene expression (**Fig. 4A**). This suggested that the lncRNA may directly influence the transcription of these genes by modulating the accessibility of transcription factor binding sites. To investigate the underlying mechanism of the profound changes in chromatin state and transcription, a transcription factor binding analysis was performed using HOMER [23]. DNA-motif enrichment analysis showed the basic region/leucine zipper motif (bZIP) to be more accessible under *LINC00607* knockout (**Fig. 4B**). Interestingly, ERG (ETS Transcription Factor ERG) and ETV2 (ETS Variant Transcription Factor 2) motifs were identified as being less accessible after *LINC00607* knockout (**Fig. 4C**). ERG and ETV2 are both members of the ETS transcription factor family, recognizing the core consensus motif GGA(A/T) [62], and are highly important for endothelial gene expression in particular [46]. Expression changes of a transcription factor might impact on the gene expression of its target gene. To exclude that the differential gene expression in response to *LINC00607* loss of function was not caused through differential expression of the transcription factors themselves, the expression of ETS family transcription factors was determined. As determined from RNA-Seq, ERG was highly expressed in normal HUVEC, whereas ETV2 expression was low. We therefore selected ERG as a candidate transcription factor mediating *LINC00607*-dependent transcription (**Fig. 4D**). Even though some of the ETS family members were differentially expressed in response to *LINC00607* knockout, the expression of ERG remained unchanged (**Fig. 4E**). These data indicate that *LINC00607*-dependent gene expression is likely mediated through changes in ERG-induced gene expression, resulting from *LINC00607*-directed changes in transcription factor binding site accessibility.

**Figure 4:**
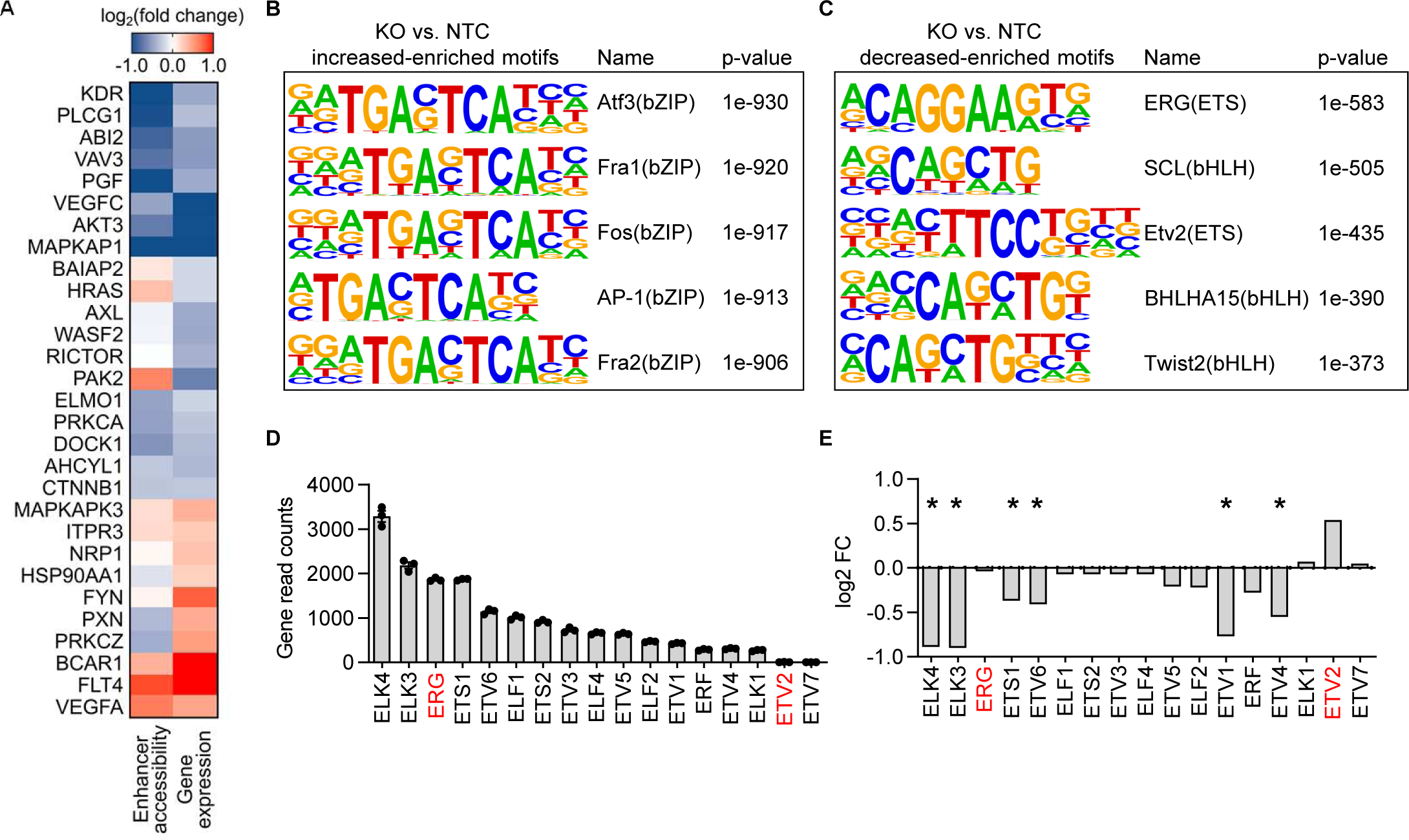
ERG drives *LINC00607*-associated gene expression. **A**, Overlap of ATAC-Seq (enhancer accessibility) and RNA-Seq (gene expression) signals after knockout of *LINC00607* in HUVEC. Indicated are genes involved in the VEGF-signaling pathway. **B-C,** HOMER DNA- motif enrichment analysis of differential accessible peaks (*LINC00607* KO vs. NTC). Five most highly increased-enriched (B) or decreased-enriched (C) transcription factor motifs are shown. **D**, Gene read counts of different transcription factors in NTC-treated HUVEC (from RNA-Seq), n=3. **E**, Mean log2 fold change (FC) of different transcription factors in the RNA-Seq comparing *LINC00607* KO and NTC control.

### LINC00607 maintains endothelial-specific chromatin states through interaction with BRG1

The changes in chromatin accessibility and to ERG-binding sites would naturally be caused by chromatin remodeling. We have previously shown that an important chromatin remodeling protein interacting with lncRNAs in endothelial cells is the SWI/SNF member BRG1 [38]. Importantly, RNA immunoprecipitation with antibodies against BRG1 yielded *LINC00607* as an interaction partner of BRG1 (**Fig. 5A**). The interaction of the lncRNA with BRG1 was specific: in contrast to β*-Actin* mRNA, *LINC00607* was not pulled down by the non-primary antibody control IgG; RNase A treatment was able to abolish the signal (**Fig. 5A**). *LINC00607* knockout did not affect *BRG1* expression (**Table S1**), which indicates *LINC00607* might influence BRG1 DNA binding activity. To test this, a lentiviral CRISPR/Cas9 knockout of *BRG1* in HUVEC was generated (**Fig. 5B&C**). Cleavage Under Targets & Release Using Nuclease (CUT&RUN) sequencing, a method to determine high-resolution mapping of DNA binding sites [58], was performed using anti-BRG1 antibodies after both non-targeting control and *BRG1* knockout in HUVEC. BRG1 binding sites were located near the transcription start sites of many genes and, upon *BRG1* knockout, these sites were lost confirming the specificity of BRG1 binding (**Fig. 5D**). To reveal the role of *LINC00607* for BRG1 binding, differentially expressed genes identified by RNA-Seq were overlapped with differential ATAC-Seq peaks having proximity to the transcriptional start site and with genes BRG1 binding sites were identified by CUT&RUN. Surprisingly, there was a strong overlap between *LINC00607* differentially regulated genes and BRG1 target genes (**Fig. 5E**). BRG1-associated genes exhibited a stronger and more significant decrease in expression after *LINC00607* knockout compared to non-BRG1-associated genes (**Fig. 5F&G**). Since the motif for the ERG transcription factor was strongly enriched in genes downregulated after *LINC00607* knockout, the described gene sets were further overlapped with a publicly available ERG Chromatin immunoprecipitation-Seq (ChIP-Seq) [29] from HUVEC. Importantly, almost all (1372 out of 1445) of the differentially accessible genes after *LINC00607* knockout overlapping with BRG1 CUT&RUN binding sites were shared with genes ERG binds close to (**Fig. 5E**).

**Figure 5:**
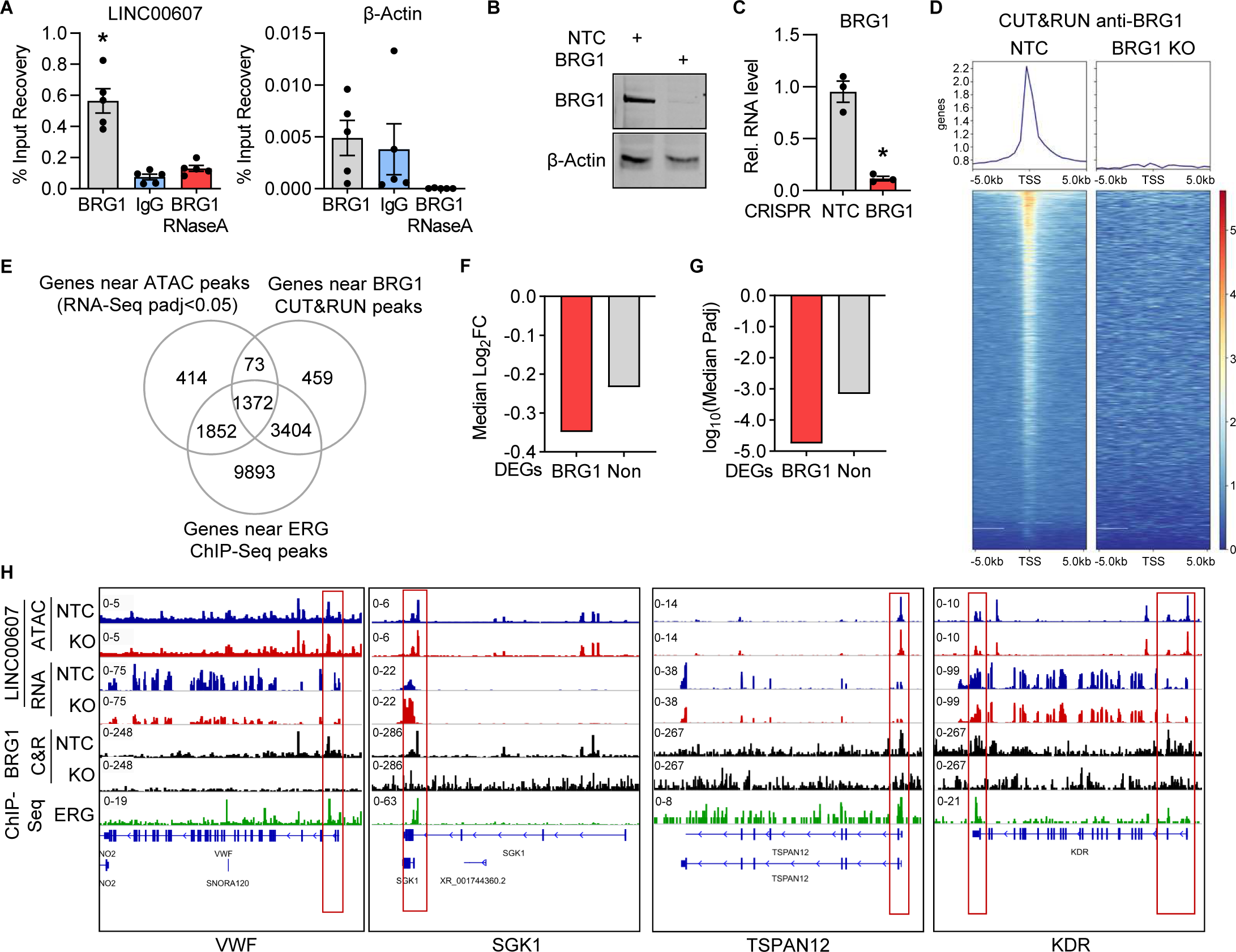
*LINC00607* functions through interaction with the chromatin remodeler BRG1. **A**, RNA-immunoprecipitation with antibodies against BRG1 with and without RNase A digestion, followed by RT-qPCR of *LINC00607* and β-*Actin*. IgG served as a non-primary antibody control. n=5. One-Way ANOVA with Bonferroni post hoc test. **B**, Western analysis with antibodies against BRG1 and β-Actin of control (NTC) or *BRG1* knockout HUVEC. **C**, RT-qPCR of *BRG1* after CRISPR/Cas9-mediated knockout, n=3. Paired t-test. **D**, Chromatin accessibility heat map of differential peaks from BRG1 CUT&RUN of control (NTC) and *BRG1* knockout (KO) HUVEC. Binding regions center-aligned to the transcription start sites (TSS) +/- 0.5 kb are shown. **E**, Venn diagram showing the overlap of genes located near a differential ATAC-Seq peak of *LINC00607* knockout, genes near a BRG1 CUT&RUN peak and genes near ERG ChIP-Seq peak. **F**, Median log2 fold change (FC) of differentially expressed genes from *LINC00607* knockout that are located near a differential ATAC-Seq peak of *LINC00607* knockout and also found near a BRG1 CUT&RUN peak (BRG1) or not (Non). **G**, Median p-adjusted value of differentially expressed genes from *LINC00607* knockout that are located near a differential ATAC-Seq peak of *LINC00607* knockout and also found near a BRG1 CUT&RUN peak (BRG1) or not (Non). **H**, Genome tracks of ATAC-Seq, RNA-Seq, BRG1 CUT&RUN and ERG ChIP-Seq. Loci of *VWF*, *SGK1*, *TSPAN12* and *KDR* are shown. ATAC-Seq and RNA-Seq of *LINC00607* KO (red) and NTC (blue) are shown. Tracks of replicates were overlaid. CUT&RUN with anti-BRG1 antibodies of NTC or *BRG1* KO HUVEC are shown in black. ChIP-Seq of ERG is shown in green.

To inspect these global associations in more detail, we checked a handful of genes highly important in endothelial cells manually: *VWF* (von Willebrand factor), *SGK1* (Serum/Glucocorticoid Regulated Kinase 1), *TSPAN12* (Tetraspanin 12) and *KDR* (Kinase Insert Domain Receptor) (**Fig 5H**). *SGK1*, *TSPAN12* and VWF were among the strongest differentially expressed genes after *LINC00607* knockout (**Fig. 3A**), and *KDR* represents a gene involved in the VEGF signaling pathway, which we described in this study to be strongly affected by *LINC00607* perturbation. Importantly, all these genes contained a BRG1 and ERG signature at their transcriptional start site (**Fig 5H**) which indicates that many LINC00607- dependent genes are also BRG1 and ERG target genes. *LINC00607* is required for the stable expression of these genes.

## Discussion

In the present study, we identified *LINC00607* to be specifically expressed in EC and to be important for vascular sprouting and regeneration. Although already constitutively highly expressed in EC, *LINC00607* itself was upregulated by hypoxia and EndMT. Through RNA- and ATAC-Seq we identified *LINC00607* as a lncRNA important for central pathways of endothelial cells, in particular for VEGF signaling. After LentiCRISPR-mediated knockout of *LINC00607,* endothelial cells exhibited an impaired response to VEGF-A in respect to vascular sprouting and a reduced ability to integrate into the vascular network of SCID mice. Mechanistically, the *trans*-acting lncRNA interacts with the chromatin remodeling protein BRG1 in order to maintain chromatin states for ERG-dependent transcription. Thereby *LINC00607* preserves endothelial gene expression patterns, which are essential for angiogenesis.

In terms of transcriptional control, lncRNAs can either act in *cis* at nearby genes, or in *trans* genome wide [54]. Overexpression of *LINC00607* restored endothelial function after *LINC00607* knockout, demonstrating that *LINC00607* acts *in trans* rather than *in cis*, because the effect of locus-disruption by CRISPR/Cas9 gene editing was overruled by the plasmid-based overexpression of *LINC00607*.

Importantly, *LINC00607* interacts with the chromatin remodeling protein BRG1. Several lncRNAs have been linked to BRG1, influencing its activity in the cardiovascular system and other tissues. For example, *EVF2* has been shown to inhibit the ATPase activity of BRG1 [10]. Additionally, lncRNAs can stabilize or destabilize BRG1 interaction with other proteins, as in the case of *MALAT1* promoting BRG1 interaction with HDAC9 [43]. lncRNAs can also affect BRG1 gene targeting. For example, we have previously shown that the lncRNA *MANTIS* guides BRG1 to specific genes related to endothelial lineage specification [38]. Our present results suggest that numerous target genes of *LINC00607*, BRG1 and ERG overlap arguing that the *LINC00607* could potentially facilitate BRG1 binding to genes linked to the endothelial phenotype. Recent studies highlight the importance of constant SWI/SNF remodeling to maintain a stable open chromatin state [24, 56]. Our present observations suggest that *LINC00607* provides a specific link for ERG securing BRG1 binding to genes maintaining the endothelial phenotype. This specific context would explain why *LINC00607* is so highly expressed in endothelial cells.

Indeed, we uncovered a large overlap between genes with altered chromatin state and differential expression after *LINC00607* knockout, and genes with binding sites for the ERG transcription factor. ERG belongs to the ETS transcription factor family, which act as key regulators of the majority of endothelial genes, as the ETS recognition motif can be found in promotors of many endothelial genes [61]. This shows the importance of *LINC00607* for the expression control of ERG-regulated genes through its interaction with BRG1. In this context, it is interesting to note that SWI/SNF is required to maintain open chromatin [24, 56]. BRG1, being the core member of SWI/SNF, could have a central role in the proposed mechanism of transcriptional control. Our findings advocate for *LINC00607* as one link between BRG1-mediated stabilization of chromatin states and ERG target gene expression in healthy endothelium.

The endothelial expression of *LINC00607* was increased by hypoxia, EndMT, and endothelial dysfunction as induced by TNFα and high glucose [11]. Under these stimuli, the upregulation of *LINC00607* matches the transcription factor binding motifs identified in the promotor analysis. Hypoxia signalling through VEGF is an important trigger for angiogenic specification of endothelial cells [52] as well as a key mechanism contributing to chronic and acute cardiovascular diseases [37]. Through the upregulation of *LINC00607* under hypoxic conditions, the pro-angiogenic endothelial phenotype could potentially be secured by tightening the interaction of *LINC00607* with BRG1. Specifically to HIF1- controlled *LINC00607* expression, we found that DMOG-dependent upregulation and acriflavine- mediated HIF inhibition altered the expression of the lncRNA. This in particular illustrates the close interaction of hypoxia-signaling, angiogenesis and *LINC00607*.

The fact that EndMT induction by TGF-β2 and IL-1β also increased the expression of *LINC00607* points towards a role for *LINC00607* in expression control beyond the endothelial phenotype. Potentially *LINC00607* guides BRG1 to genes involved in EndMT. In line with this, *LINC00607* is also expressed in certain malignant cells. In this context, *LINC00607* was upregulated in doxorubicin-resistant thyroid cancer cells [41]. Furthermore, *LINC00607* was described to be required for tumor proliferation of osteosarcoma cells [68] and was downregulated in lung adenocarcinoma [67]. Linking these findings to the present study, it could be speculated that under basal conditions, *LINC00607* guides BRG1 to ERG target genes and during endothelial dysfunction to pro-angiogenic genes to maintain an open and accessible chromatin state for ERG.

Taken together, these findings suggest that *LINC00607* is involved in securing endothelial BRG1 and ERG-dependent target gene expression, as well as appropriate responses to stress and cardiovascular diseases. The function of *LINC00607* in cardiovascular diseases with hypoxia signaling needs more investigation as *LINC00607* might be a potential therapeutic target or clinical marker.

## Acknowledgments

We thank Katalin Pálfi for excellent technical assistance and Dominik Fuhrmann for help with the hypoxia chamber.

## Author contributions

FB, JAO, TW, SK, FJM, RPB and MSL designed the experiments. FB, JAO, TW, SG, JIP, GB, TL, SS, SH, SK, FJM and MSL performed the experiments. FB, JAO, TW, SG, GB and MSL analyzed the data. FB, TW, SG and MHS performed bioinformatics. AHB and RAB helped with research design and advice. FB, RBP and MSL wrote the manuscript. All authors interpreted the data and approved the manuscript.

---

**Sup. Fig. 1:**
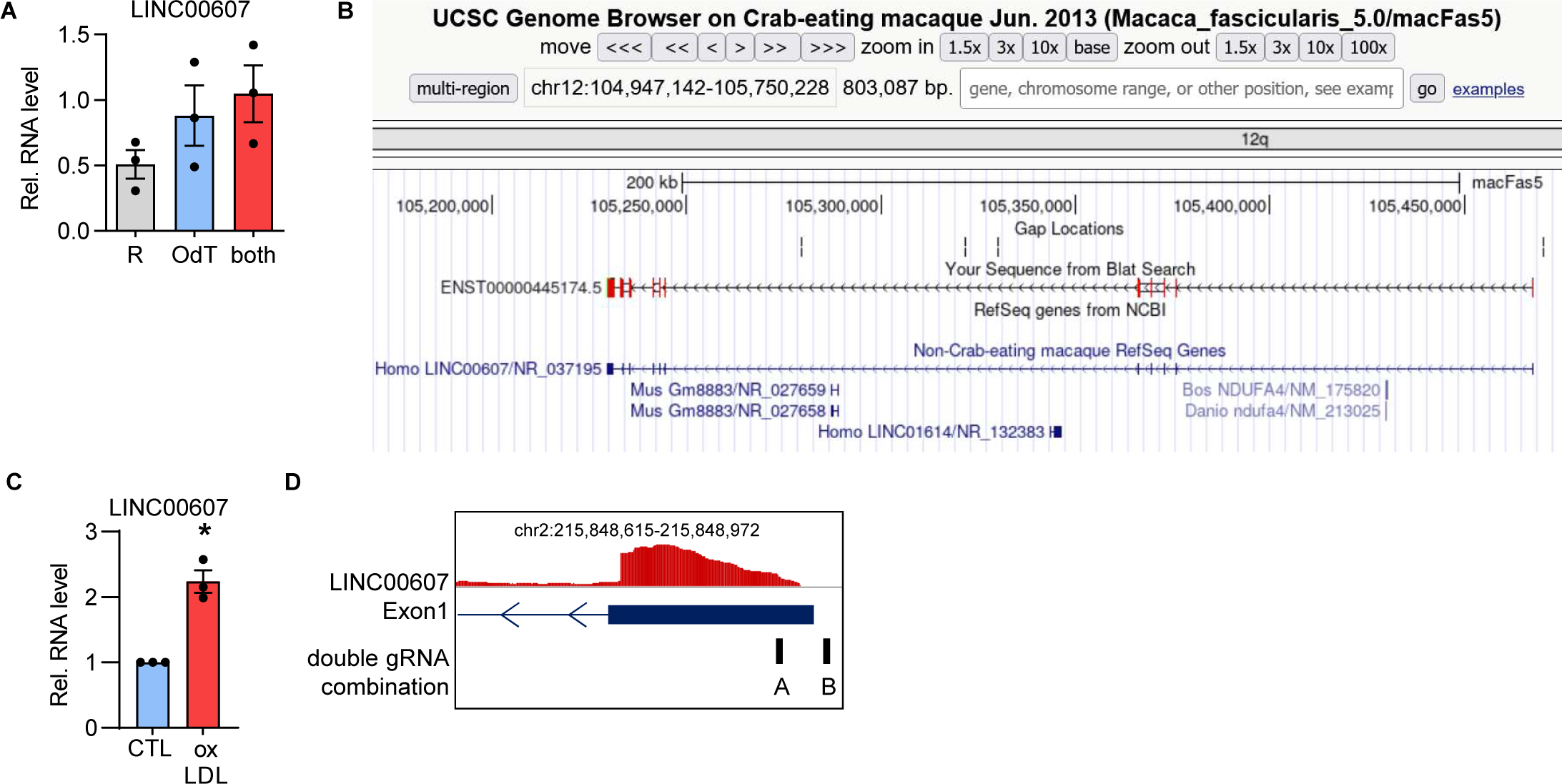
**A**, Relative RNA expression of *LINC00607* in HUVEC after RT-qPCR with either only random hexamer primers (R), only Oligo(dT) primers (OdT) or the combination of both (both). n=3, Paired t-test. **B**, UCSC Genome browser view of the *LINC00607* homologue in *Macaca fascicularis*. BLAT nucleotide sequence alignments of cDNA of human versus *Macaca fascicularis* are shown in red. **C**, Relative *LINC00607* expression in HUVEC treated with oxLDL (16h, 10 µg/mL). n=3, Paired t-test. **D**, Scheme of gRNAs used for CRISPR/Cas9-mediated KO of *LINC00607*. Error bars are defined as mean +/- SEM. *p<0.05.

**Sup. Fig. 2:**
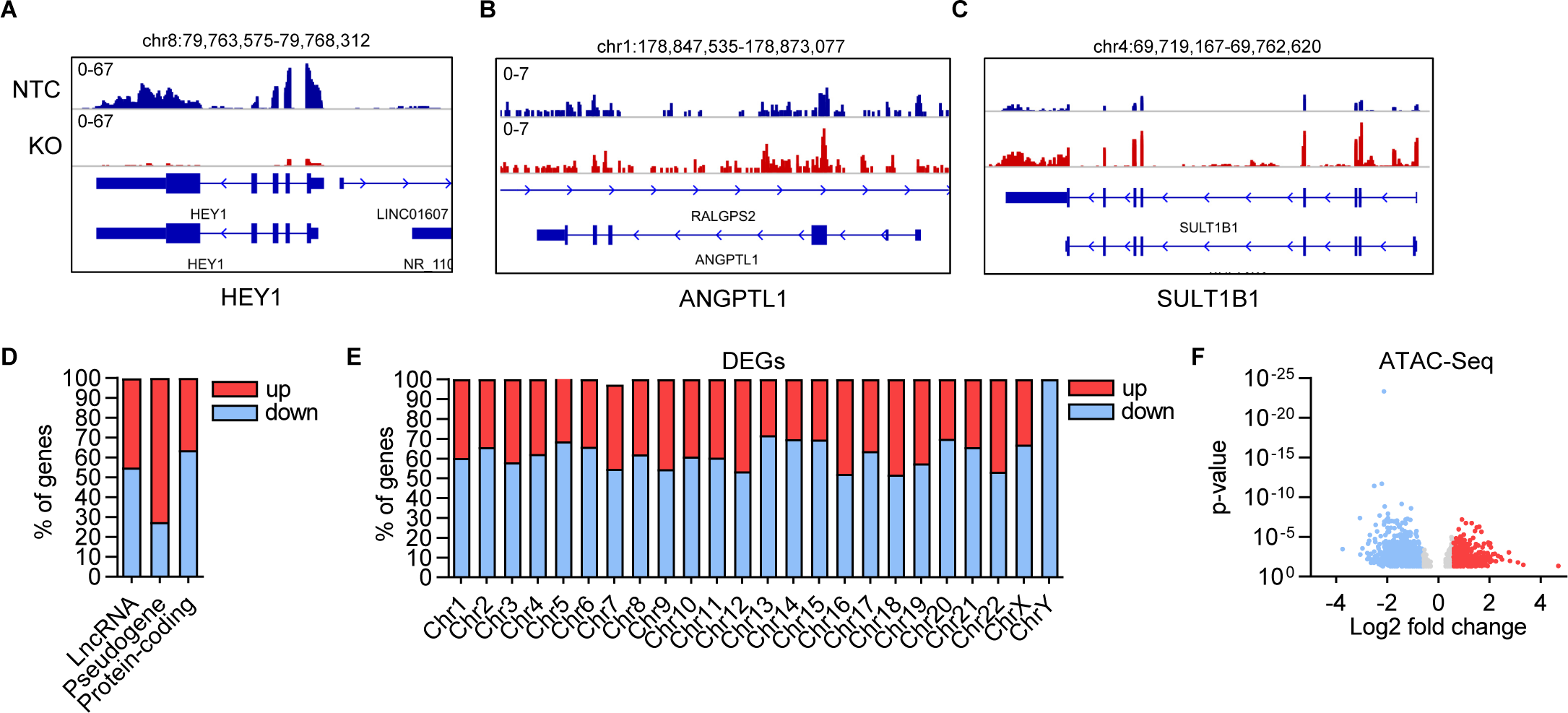
**A-C**, Examples of significantly down- and up-regulated genes after *LINC00607* KO. IGV genome tracks of the *HEY1*, *ANGPTL1* and *SULT1B1* locus. Shown are genomic tracks of *LINC00607* KO (red) and NTC (blue); tracks of three replicates are overlaid. **D-E,** Percentage of genes belonging to lncRNAs, pseudogenes or protein-coding genes (D) or chromosomal distribution and percentage of genes (E) up- or downregulated (DEGs) in the RNA-Seq after *LINC00607* KO in HUVEC. Only genes with a log2 fold change of +/-0.585, a basemean expression of 5 and a p-adjusted value <0.05 are shown. Genomic coordinates correspond to hg38. **F**, Volcano plot of ATAC-Seq showing the log2 fold change (KO vs. NTC) of all peaks against their p-value.

